# Dynamic genome-scale modeling of *Saccharomyces cerevisiae* unravels mechanisms for ester formation during alcoholic fermentation

**DOI:** 10.1101/2022.05.27.493771

**Authors:** William T. Scott, David Henriques, Eddy J. Smid, Richard A. Notebaart, Eva Balsa-Canto

## Abstract

Fermentation employing *Saccharomyces cerevisiae* has produced alcoholic beverages and bread for millennia. More recently, *S. cerevisiae* has been used to manufacture specific metabolites for the food, pharmaceutical, and cosmetic industries. Among the most important of these metabolites are compounds associated with desirable aromas and flavors, including higher alcohols and esters. Although the physiology of yeast has been well-studied, its metabolic modulation leading to aroma production in relevant industrial scenarios such as winemaking is still unclear. Here we ask what are the underlying metabolic mechanisms that explain the conserved and varying behavior of different yeasts regarding aroma formation under enological conditions? We employed dynamic flux balance analysis (dFBA) to answer this key question using the latest genome-scale metabolic model (GEM) of *S. cerevisiae*. The model revealed several conserved mechanisms among wine yeasts, e.g., acetate ester formation is dependent on intracellular metabolic acetyl-CoA/CoA levels, and the formation of ethyl esters facilitates the removal of toxic fatty acids from cells using CoA. Species-specific mechanisms were also found, such as a preference for the shikimate pathway leading to more 2-phenylethanol production in the Opale strain as well as strain behavior varying notably during the carbohydrate accumulation phase and carbohydrate accumulation inducing redox restrictions during a later cell growth phase for strain Uvaferm. In conclusion, our new metabolic model of yeast under enological conditions revealed key metabolic mechanisms in wine yeasts, which will aid future research strategies to optimize their behavior in industrial settings.

## 1 INTRODUCTION

The genetics and metabolism of the yeast, *Saccharomyces cerevisiae*, have been studied extensively as a model eukaryotic organism (Botstein et al., 1997, Liti, 2015). However, despite its general use for industrial processes such as winemaking, variability in fermentation performance, especially in growth and aroma production, is not entirely understood (Hirst and Richter, 2016). Since prominent aromas, e.g., higher alcohols and esters, are partially produced during active growth, factors that affect yeast growth simultaneously influence essential aroma formation (Dekoninck, 2012). Furthermore, commercial yeast strains vary by orders of magnitude in their aroma production (Steensels et al., 2014, Gonzalez and Morales, 2017, Pérez et al., 2021). This variation has been associated with a myriad of genetic (Peter et al., 2018) and environmental factors such as nitrogen and micronutrients concentration levels (Su et al., 2021), temperature (Rollero et al., 2015), pH (Lam et al., 2014), ethanol concentration levels (Snoek et al., 2016), and the presence of toxins (Viegas et al., 1989).

Commercial strains of wine yeast respond to these factors differently in poorly understood ways, but their responses appear to be correlated with specific parts of yeast metabolism, such as those involved in lipid formation and membrane composition (Henderson et al., 2013). Moreover, it has been suggested that particular aromas are synthesized because of the detoxification of medium-chain fatty acids. It is also speculated that aroma formation is part of a metabolic process used to balance the acetyl-CoA/CoA ratio (Mason and Dufour, 2000). Understanding the differences in metabolism in yeast strains will lead to a greater ability to control and manipulate aroma-related performance. That greater metabolic understanding will facilitate insight into the specific mechanisms inducing aroma formation.

Several studies have examined the production of volatile organic compounds (VOC) or aroma (Miller et al., 2007, Carrau et al., 2008, Seguinot et al., 2018). Some of these studies have combined omics approaches, exploring the link between yeast gene expression and metabolomics (Dunn et al., 2005, Minebois et al., 2020) or analyzing the transcriptome and metabolome profiles of yeast strains to assess the expressed genes on aroma formation during wine fermentation (Rossouw et al., 2008). Although there is experimental evidence for differences in the metabolism of various strains and genes involved in aroma formation, these experimentally-derived large data sets can be challenging to generate and analyze, especially in terms of finding the most important differences relevant to the metabolism being studied. As an alternative, mathematical modeling of yeast metabolism can provide a more comprehensive means to examine how yeast metabolism changes during the entire course of wine fermentation and which parts of overall metabolism are interrelated.

Metabolic modeling of yeast under enological conditions has been reported for some time (Boulton, 1980, Cramer et al., 2002, Sainz et al., 2003, Pizarro et al., 2007). The general approach has been to use Flux Balance Analysis (FBA) with a subset of metabolites and metabolic reactions found in yeast during alcoholic fermentation (Quiros et al., 2013), often augmented with dynamic FBA (dFBA) to account for the time-dependent nature of wine fermentations and their associated yeast metabolism. More recently, the number of metabolites and reactions included in models of *Saccharomyces cerevisiae* has grown to be more comprehensive to the point where they are often referred to as genome-scale metabolic models (GEMs).

Consensus GEMs for yeast have developed from model *iFF708* (1172 reactions) (Forster et al., 2003) to increasingly comprehensive models (Nookaew et al., 2008, Aung et al., 2013) (1413 and 3498 reactions, respectively), with the newest consensus model being *Yeast8* (3498 reactions) (Lu et al., 2019). This recent model has been further curated to include some lipid synthesis pathways and amino acid degradation pathways to account for the production of higher alcohols, carboxylic, acids and esters (Scott et al., 2020), which have been proven to be key in imparting desirable aromas to alcoholic beverages (for more details, see review by (Dzialo et al., 2017)).

Vargas et al. (2011) proposed the application of dFBA to less extensive GEMs for studying wine fermentations (Vargas et al., 2011); however, that study lacked exploration into intracellular flux behavior and insight into how kinetic constraints impact prediction performance. Jouhten and coworkers used dFBA with *Yeast5* (Herrgård et al., 2008) to simulate the dynamic metabolic response of *S. cerevisiae* exposed to sudden oxygen depletion (Jouhten et al., 2012). Also, Sanchez and coworkers devised a dFBA framework with *Yeast5* to successfully calibrate different types of data from aerobic fed-batch and anaerobic batch cultivations (Sánchez et al., 2014). These two studies nevertheless pertained to *S. cerevisiae* grown under glucose-limited conditions while nitrogen-limited growth of *S. cerevisiae* is relevant to enological conditions.

Yeast8 has been combined with constraint-based modeling approaches to understand yeast metabolism in a few studies. However, the use of this yeast GEM has thus far been confined to glucose-limited, aerobic conditions (Moreno-Paz et al., 2022), nutrient-rich cases (Henriques et al., 2021b) and/or under non-transient model scenarios (Scott et al., 2021b), thus limiting its scope and applicability to accurately predict enological conditions.

In this study, the *Yeast8.5.0* GEM (Lu et al., 2019), along with the dFBA framework proposed by (Henriques et al., 2021b), were employed to predict the metabolic behavior of *Saccharomyces cerevisiae* commercial strains under enological conditions. The dFBA framework proposed by Henriques et al. has advantages versus previous dFBA frameworks using yeast GEMs because it contains a multiphase multiobjective implementation of a parsimonious flux balance analysis (pFBA) and a method for capturing changes in the biomass equation at different growth phases. Our model framework was mainly used to model nitrogen-limited, anaerobic growth of *S. cerevisiae* with appropriate kinetic constraints and utilizing a biomass equation tailored for enological conditions. After calibrating the model with published experimental data from four commercial yeast strains (Scott et al., 2021a), the goodness of fit of the model was assessed for each strain to evaluate the accuracy and quality of the model simulations. We analyzed the metabolic flux distributions corresponding to the best fit to the experimental data. Our analysis revealed conserved behavior across strains. However, some strain-specific mechanisms were also noticed, particularly in the Uvaferm and Opale strains.

## 2 EXPERIMENTAL PROCEDURES

### 2.1 Experimental data

The experimental data for this study come from a previously described study published by Scott and coworkers (Scott et al., 2021a). The yeast strains used in experiments were Uvaferm 43™(Uvaferm), Lalvin R2™ (R2), Lalvin ICV Opale™ (Opale), and Vitilevure™ Elixir Yseo (Elixir), which are all commercial strains developed by Lallemand (Lallemand, Montreal, Quebec). All yeast strains were acquired from the UC Davis Enology Culture Collection. Fermentations were carried out for each strain in MMM synthetic grape juice medium under nitrogen-limited conditions (yeast assimilable nitrogen, YAN ~ 120 mg/L). More specifically, sugar (220 g/liter; ~ 22.0oBrix), as a 1:1 mixture of glucose (110 g/liter) and fructose (110 g/liter), MMM synthetic grape juice medium was prepared according to the method of Giudici and Kunkee (1994), as previously reported (Henderson et al. 2011). Regarding YAN, to make one liter of MMM medium, 0.1g, of ammonium phosphate, 0.1g L-tryptophan, 0.2g L-arginine*HCl, 1.0g L-proline, and 2 g of vitamin-free Casamino acid were added. From analysis reported in Nolan (1971), 2g of Vitamin-Casamino acid (Difco) consists of 0.649 mM, lysine, 0.194 mM histidine, 0.709 mM ammonia, 0.227 mM arginine, 0.029 mM cysteic acid, 0.657 mM asprartic acid, 0.417 mM threonine, 0.623 mM serine, 1.604 mM glutamic acid, 1.237 mM proline, 0.311 mM, glycine, 0.440 mM alanine, 0.600 mM valine, 0.225 mM methionine, 0.392 mM isoleucine, 0.877 mM leucine, 0.007 mM tyrosine, and 0.091 mM phenylalanine (Nolan 1971). The medium also contained 1000 mg/L of proline which is not commonly used by *Saccharomyces cerevisiae* as a source of YAN under enological conditions. The pH of the MMM medium was adjusted to ~ 3.25 using 3 N KOH. All the experiments were carried out at a constant temperature of 20 °C. These yeast strains were selected based on their different fermentation and aroma-producing performance attributes. Furthermore, the manufacturers present qualitative characterizations of these strains, but publish information that lacks quantitative distinctions (see Table 1). For additional information pertaining to the media preparation, yeast strains, culture conditions, or fermentation sampling that were performed to generate the experimental data used in this study, see report from Scott and coworkers (Scott et al., 2021a).

**Table 1.**
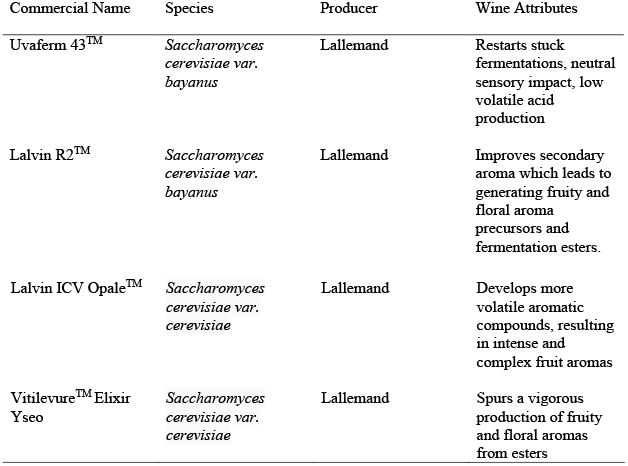
Main enological properties of select yeast strains

| Commercial Name | Species | Producer | Wine Attributes |
| --- | --- | --- | --- |
| Uvaferm 43<sup>TM</sup> | *Saccharomyces cerevisiae* var. *bayanus* | Lallemand | Restarts stuck fermentations, neutral sensory impact, low volatile acid production |
| Lalvin R2<sup>TM</sup> | *Saccharomyces cerevisiae* var. *bayanus* | Lallemand | Improves secondary aroma which leads to generating fruity and floral aroma precursors and fermentation esters. |
| Lalvin ICV Opale<sup>TM</sup> | *Saccharomyces cerevisiae* var. *cerevisiae* | Lallemand | Develops more volatile aromatic compounds, resulting in intense and complex fruit aromas |
| Vitilevure<sup>TM</sup> Elixir Yseo | *Saccharomyces cerevisiae* var. *cerevisiae* | Lallemand | Spurs a vigorous production of fruity and floral aromas from esters |

### 2.2 Genome-scale metabolic model (GEM)

The GEM used in this study was *Yeast8.5.0* (Lu et al., 2019), publicly available on GitHub (see link: https://github.com/SysBioChalmers/yeast-GEM). The GEM contains 2742 metabolites, 4058 reactions, and 1150 genes. The GEM was initially developed for *S. cerevisiae* S288C, a haploid laboratory strain that is not employed in winemaking. However, since this study was applied to fermentations under enological conditions, we modified the GEM to appropriately reflect the anaerobic state of metabolism. In our approach, we proceeded as suggested by Heavner and coworkers (Heavner et al., 2013), constraining 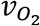 to zero (LB=UB=0 [mmol/g DW h]), allowing unrestricted uptake of ergosterol (r_1757), lanosterol (r_1915), zymosterol (r_2106), 14-demethyllanosterol (r_2134), and ergosta-5,7,22,24(28)-tetraen-3beta-ol (r_2137) and oleate (r_2189). In addition, pathways including the oxaloacetate-malate shuttle and glycerol dehydrogenase reaction were unrestricted as described by Sanchez and coworkers (Sánchez et al., 2017, Sánchez et al., 2019) (in the model, this was achieved by blocking reactions r_0713, r_0714, and r_0487). Heme A was also removed since it is not used under anaerobic conditions. All the changes related to anaerobiosis were applied either at the beginning of the fermentation or after the depletion of the initially dissolved oxygen in the simulated media. Moreover, *Yeast8.5.0* includes expanded coverage of aroma-associated pathways such as an extended Ehrlich pathway, more ester formation reactions, and enhanced sulfur reduction pathways as previously performed and described in the literature (Scott et al., 2020).

The GEM used in this study differs from that used in the (Henriques et al., 2021b) study. Henriques et al. expanded *Yeast8 (v.8.3*.1) to include 38 metabolites and 50 reactions to explain secondary metabolism. Furthermore, 13 aroma-impact molecules were added to GEM (see Table S1 in (Henriques et al., 2021b) for more details). The GEM used in this work is *Yeast8 (v.8.5.0*). This version of *Yeast8* contains at least 72 reactions and 50 metabolites related to amino acid degradation and ester synthesis, which induce the formation of some aroma compounds. Moreover, *Yeast 8.5.0* expanded coverage is based on the work of (Scott et al., 2020) (see version release details for the complete list of updates here: https://github.com/SysBioChalmers/yeast-GEM/releases).

### 2.3 Flux balance analysis

Flux balance analysis (FBA) (Varma and Palsson, 1994, Orth et al., 2010) is a modeling framework based on knowledge of reaction stoichiometry and mass/charge balances. The framework relies on the pseudo-steady-state assumption (no intracellular accumulation of metabolites occurs). The well-known expression captures FBA:

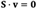

where **S** is the stoichiometric matrix of (*n* metabolites by *m* reactions), and **b** is a vector of metabolic fluxes. The number of unknown fluxes is higher than the number of equations; thus, the system is undetermined. Still, it is possible to find a unique solution under the assumption that cell metabolism evolves to pursue a predetermined goal which is defined as the maximization (or minimization) of a particular objective function (J):

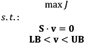

where **LB** and **UB** correspond to the lower and upper bounds on the estimated fluxes. Examples of objective functions J include growth rate, ATP, or the negative of nutrient consumption, etc.

The solution to an FBA problem is often not unique, i.e., several different combinations of fluxes may lead to the same optimum. A common approach to finding a single solution is to further constrain the problem with prior knowledge or biological intuitions. Often, the so-called parsimonious FBA (pFBA) is used. The underlying idea is to find the solution that minimizes overall flux through the metabolic network (a proxy for minimizing the total necessary enzyme mass to implement the optimal solution). (Lewis et al., 2010, Machado et al., 2014).

### 2.4 Parameter estimation

Parameter estimation aims to compute the unknown parameters - growth-related constants and kinetic parameters - that minimize some measure of the distance between the data and the model predictions. The maximum-likelihood principle yields an appropriate measure of such a distance (Walter and Pronzato, 1997):

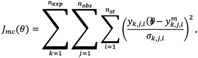

where *n_exp_*, *n_obs_* and *n_st_ σ_k,j,i_*, 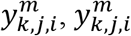 and 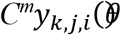 corresponds to model predicted values, *X* and *C*. Observation functions were included for CFUs and OD_600_ to scale viable cell mass (*X*_V_) and active cell mass (*X*_A_), respectively.

Parameters are estimated by solving a nonlinear optimization problem where the aim is to find the unknown parameter values (*θ*) to minimize 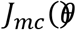 subject to the system dynamics - the model- and parameter bounds (Balsa-Canto et al., 2010).

To assess the confidence in the parameter estimates we estimated the covariance matrix from the Fisher Information Matrix:

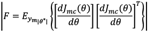

where E represents expected value and θ* is the optimal parameter value. Remark that the evaluation of the derivatives of the log-likelihood function (J_mc_) with respect to the parameters, requires the computation of the sensitivities of the measured states with respect to the parameters. The Cramèr-Rao inequality provides a lower bound on the covariance of the estimators:

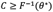

The confidence interval for a given parameter is then given by:

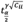

where t^γ^_α/2_ is given by the Students t-distribution, γ regards the number of degrees of freedom and α is the (1-α) 100% confidence interval.

### 2.5 The Multi-Phase Multi-Objective Flux Balance Analysis Framework

In this work, we have adapted the model described in (Henriques et al., 2021b) that accounts for the dynamic nature of batch fermentation and divides the process into five phases: the lag phase, exponential growth, growth under nitrogen limitation, stationary phase, and decay. Each phase is characterized by a cellular objective and a set of constraints. Their duration is imposed by the estimated parameters TL, TCA, TS, and TD. Interested readers can find a detailed description of the model equations for the different phases (Henriques et al., 2021b). Here we summarize some of the most relevant characteristics for each phase:

- Lag phase: The objective is to maximize ATPase expenditure (equivalent to maximizing ATP production).
- Exponential growth phase (until the nitrogen sources were nearly exhausted): The objective is to maximize the growth rate.
- Carbohydrate accumulation phase: The objective is to maximize protein production with constrained carbohydrate accumulation.
- Stationary and decay phases: The objective is to maximize ATP and protein production.

These objectives were chosen assuming that cells behave optimally throughout the process. Typically, the cellullar objective is assumed to be maximizing biomass. However, in batch conditions, it is observed that growth is not possible in specific phases. In those phases we assume that cells will behave efficiently to conserve existing biomass. In particular, we chose protein and ATP maximization objectives during the carbohydrate, stationary, and decay phases. The rationale behind is that yeast cells are subject to protein turnover and when it is not possible to sustain growth, yeast cells will try to retain as much nitrogen as possible. Further, enforcing maximum ATP production, we assume that metabolism will be as energy efficient as possible.

Constraints on the external fluxes were modeled using kinetic models in ordinary differential equations. The transport of hexoses was described using Michaelis-Menten-type kinetics subject to non-competitive inhibition of ethanol (Hjersted et al., 2007). The production of ethanol, the highest alcohols, carboxylic acids, and esters, was proportional to the amount of hexoses transported. The transport of nitrogen sources was encoded with Michaelis-Menten-type kinetics for the case of ammonia and mass action for the amino acids.

During the exponential phase, the protein content on the biomass was assumed to be 50%. The level of mRNA is assumed to be proportional to the protein content. Maintenance of growth-associated ATP (GAM) was also updated to account for the polymerization costs of the different macromolecules (proteins, RNA, DNA, and carbohydrates):

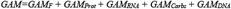

where *GAM_F_* is a species or strain-dependent parameter estimated from the data, and the rest are the polymerization costs of the different biomass precursors (adapted from (Lu et al., 2019)). All scripts necessary to reproduce the results are distributed (https://sites.google.com/site/amigo2toolbox/examples).

### 2.6 Analysis of dynamic metabolic fluxes

Here, we evaluated metabolic pathways using a flux ratio, which measures net flux over time during growth and stationary phases. In particular, we computed the integral of each flux multiplied by the biomass (mmol . h ^−1^) over time and normalized its value with the accumulated flux of consumed hexoses (glucose and fructose):

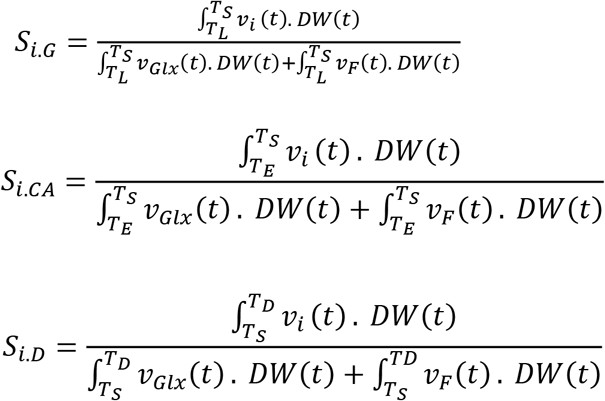

where *S_i,G_* corresponds to the flux score *i* during growth, *S_i,CA_* corresponds to the score during stationary and *S_i,D_* decay phases, *v_i_(t)* (mmol . h^−1^ . DW^−1^) is the flux under scrutiny, v_Glx(t)_ (mmol . h^−1^ . DW^−1^) is the glucose flux, *v_Fr(t)_* (mmol . h^−1^ . DW^−1^) is the flux of fructose and DW is the predicted dry weight biomass (g). The results correspond to the mmol of compound produced per mmol of hexose consumed × 100 (denoted mmol / mmolH). Score values indicate the overall impact of each reaction in the net oxidation or reduction of electron carriers during the given phase of fermentation.

### 2.7 Computing Environment

The modeling was performed in MATLAB^®^ 2020b (The MathWorks, Inc., Cambridge, MA, USA) using Cobra Toolbox 3.0 (Heirendt et al., 2019) and implemented on a Windows 10 (Microsoft Corporation, Redmond, WA, USA) Intel^®^ (Intel Corporation, Santa Clara, CA, USA) Core™ i7-7500 CPU @ 2.70 GHz–2.90 GHz processor. The GEM was imported into MATLAB as an SBML file and curated using Cobra Toolbox. Git version 2.3.0 was installed before cloning COBRA with GitHub and initializing COBRA in MATLAB.

The multi-phase multi-objective genome-scale model framework was implemented as a script for AMIGO2 toolbox (Balsa-Canto et al., 2016) to facilitate parameter estimation, simulation, and quality of fit analyses. From the possibilities available in AMIGO2, we selected to solve the model using the initial value problem solver CVODES (Hindmarsh et al., 2005) and a Nelder-Mead method (fminsearch, in MATLAB) to optimize parameter values in reasonable computational time. In addition, the tool automatically computes the FIM-based confidence intervals for the optimal parameter values.

## 3 RESULTS AND DISCUSSION

### 3.1 Multi-phase multi-objective dynamic flux balance analysis framework

We adapted the model reported earlier (Henriques, et al., 2021) to describe the intracellular fluxes. Here, we divided the process into five phases in which cellular objectives and flux constraints must be adapted: lag, exponential growth, carbohydrate accumulation, stationary, and decay (see Fig. 1). The duration of the phases was determined by the parameters t_L_, t_E_, t_s_, and t_D_.

**Figure 1.**
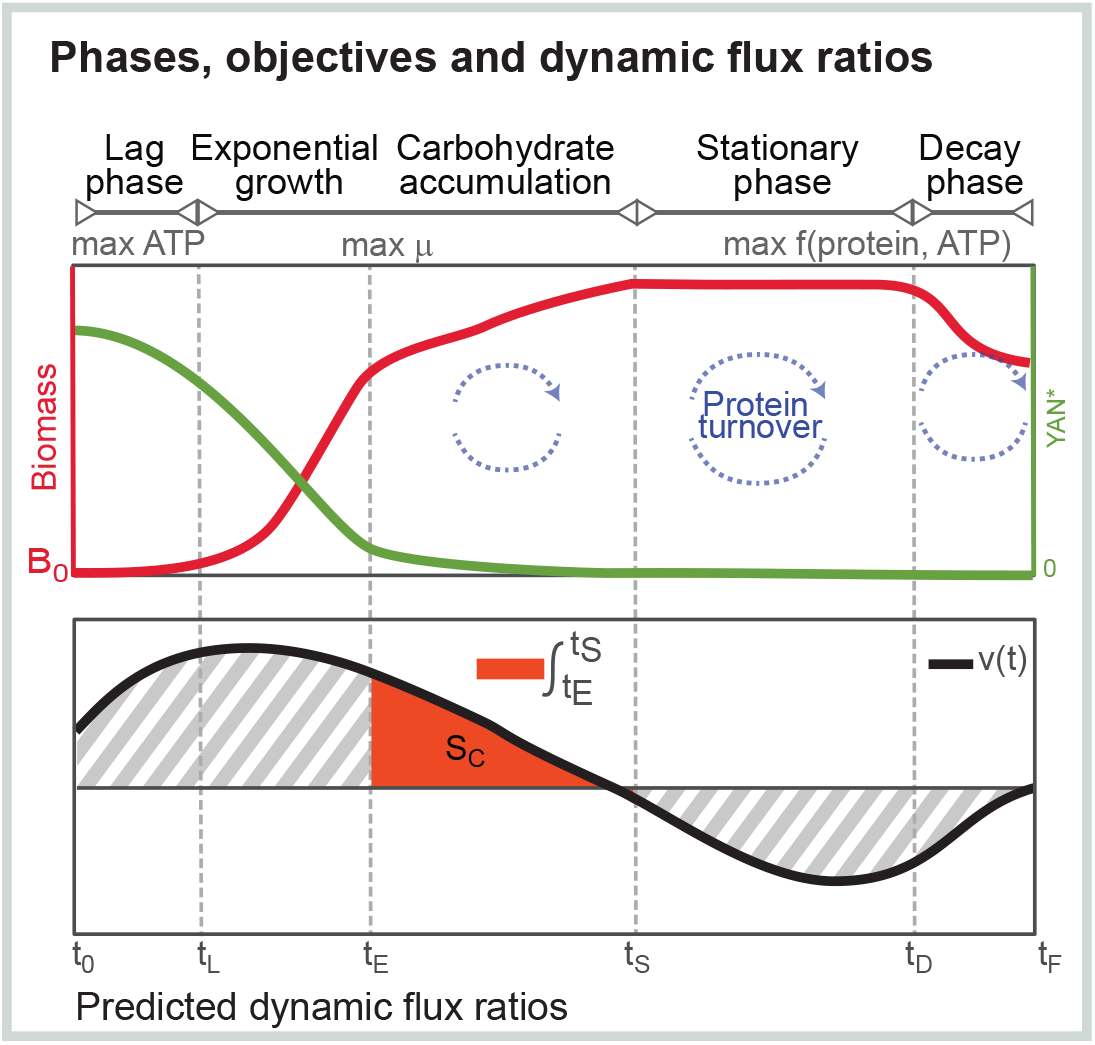
Details on the implementation of the multi-phase and multi-objective dynamic genome-scale model to simulate batch fermentation. Multi-phase and multi-objective dynamic FBA and methodology to compute dynamic flux rates. The process starts at t_0 ¼ 0_ and ends at t_F_; the timing of each phase t_L_, t_E_, t_S_, and t_D_ is computed through parameter estimation.

In its original version, the model relied on a dynamic biomass equation dependent on the amount of YAN in the medium. Nevertheless, this equation required estimating several parameters without information about biomass composition (protein, mRNA, carbohydrates). In addition, the model could not accurately explain the observed behavior of the non-*cerevisiae* species (Uvaferm). This strain showed a long growth period with virtually no nitrogen sources available. To address this, we replaced the phase regarded as growth under nitrogen limitation with a phase characterized by carbohydrate accumulation (see (Henriques et al., 2021a) for further details).

Most nitrogen sources, except glycine, were consumed primarily during the exponential growth phase. In this period, the cellular objective was the maximization of biomass. In contrast to the previous model, during carbohydrate accumulation, we assumed cells maximize protein and activated the procedure to simulate protein turnover (described in (Henriques et al., 2021b)). Also, to simulate carbohydrate accumulation during this period, an exchange flux for this compound (s_3717[c]) was added to the stoichiometric network determined by the equation:

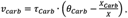

where *X_Carb_*(g/L) is the carbohydrate quantity present in the biomass, *X* is the biomass (g/L), *θ_Carb_* is the final carbohydrate content and *τ_Carb_* is the parameter controlling the convergence rate towards *θ_Carb_* (see supplementary code example (dynamic genome-scale modeling of yeast fermentation): https://sites.google.com/site/amigo2toolbox/examples).

Higher alcohol production can occur due to the assimilation/catabolism of amino acids or due to *de novo* synthesis. In its previous version, the model considered that higher alcohol production started during growth under nitrogen limitation and was prolonged through the stationary phase. The experiments considered in that work included ammonium diphosphate supplementation, and the final concentrations of higher alcohols were higher than those found in the present work. Here, the relative contributions of higher alcohols produced during the first hours of growth were relevant. Thus, to account for this effect, during exponential growth, we set the production of isoamyl alcohol, 2-phenyl ethanol, and isobutanol at the same flux as the consumption of the corresponding amino acids (leucine, phenylalanine, and isoleucine).

### 3.2 Goodness of fit of the model against enological fermentation data

This study employed data from experimental fermentations carried out under enological conditions (see (Scott et al., 2021a)) to calibrate model parameters. The final model contained 50 ODEs, consisting of 60 parameters determined from time-course data for all measured extracellular metabolites and biomass. The best fit to the data for glucose and fructose uptake, and the secretion of ethanol, acetate, malate, succinate, and glycerol, are shown in Fig. 2 for the four strains. For the primary metabolites, glucose in the medium was consumed first, followed by fructose since hexose transporters in the cytoplasm have a higher affinity for the former (Fig. 2). A stoichiometrically accurate yield of ethanol (~101 g/L) was predicted for the strains, reaching an average simulated concentration of 100 g/L with the initial starting concentration of total sugar used in this study (~220 g/L) (Fig. 2). Moreover, the values were within the range of similar enological fermentations (Ribéreau-Gayon et al., 2000). Various other essential extracellular metabolites are also measured and predicted throughout fermentation (Fig. 2), including glycerol, malate, succinate, and acetate, with average yields among strains of ~7 g/L, ~3.2 g/L, ~1.4 g/L, and ~0.5 g/L, respectively.

**Figure 2.**
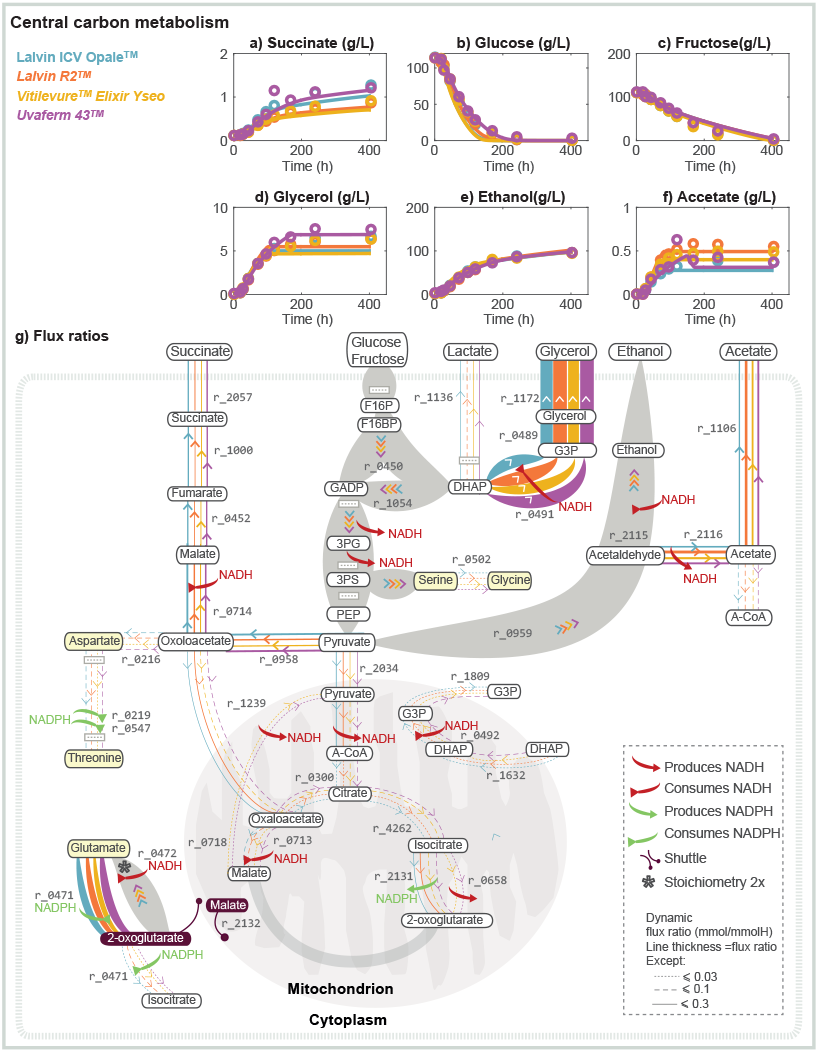
Overview of central carbon metabolism. Panels a to f depict model predictions versus the experimental data on extracellular metabolite concentrations associated with glycolysis and central carbon metabolism for the four strains. Panel g illustrates the predicted intracellular dynamic flux ratios during the carbohydrate accumulation phase, showing how the four commercial strains employ different redox balance strategies. These differences result in the differential production of relevant external metabolites such as succinate (a), glycerol (d), ethanol (e), and acetate (f).

Consumption of glucose and fructose, and production of ethanol simulations were representative of the enological conditions achieving excellent fits against the experimental dynamic profiles (with median R^2^ values of 0.98, 0.96, and 0.98 among the strains) (see Fig. 2 and Table S1). YAN and biomass (dry cell weight) curves were also successfully predicted for all the strains, where the median R^2^ values among the strains are 0.92 and 0.96, respectively (see Table S1). In addition, the production of some essential by-products of wine fermentation, including glycerol, succinate, and acetate, were successfully simulated by the model achieving moderately good fits among the strains with median R^2^ values of 0.89, 0.81, and 0.77 (Fig. 2 and Table S1). Maximum glycerol (~6.9 g/L) and succinate (~1.3 g/L) produced during the fermentation were quantitatively well predicted using the model, while acetate (~0.5 g/L) prediction was reasonably good (Fig. 2).

The nitrogen consumed by the yeast was in the form of amino acids and ammonium in the MMM synthetic grape medium, and the model profiles for the nitrogen sources for the four strains are shown (Fig. 3). As expected, since it has been well documented that wine fermentations are nitrogen-limited as suggested in several studies (Ingledew and Kunkee, 1985, Cramer et al., 2002, Varela et al., 2004), the yeast assimilable nitrogen (YAN) (a measure of the concentration of all nitrogen in the free amino acids and ammonium) was consumed very rapidly. At that point, the maximum cell density was reached. Figures 2, 3, and 4 showed a mixture of AA and NH3. The dFBA also accurately simulated the profiles of several important VOCs for the four strains (Fig. 4).

**Figure 3.**
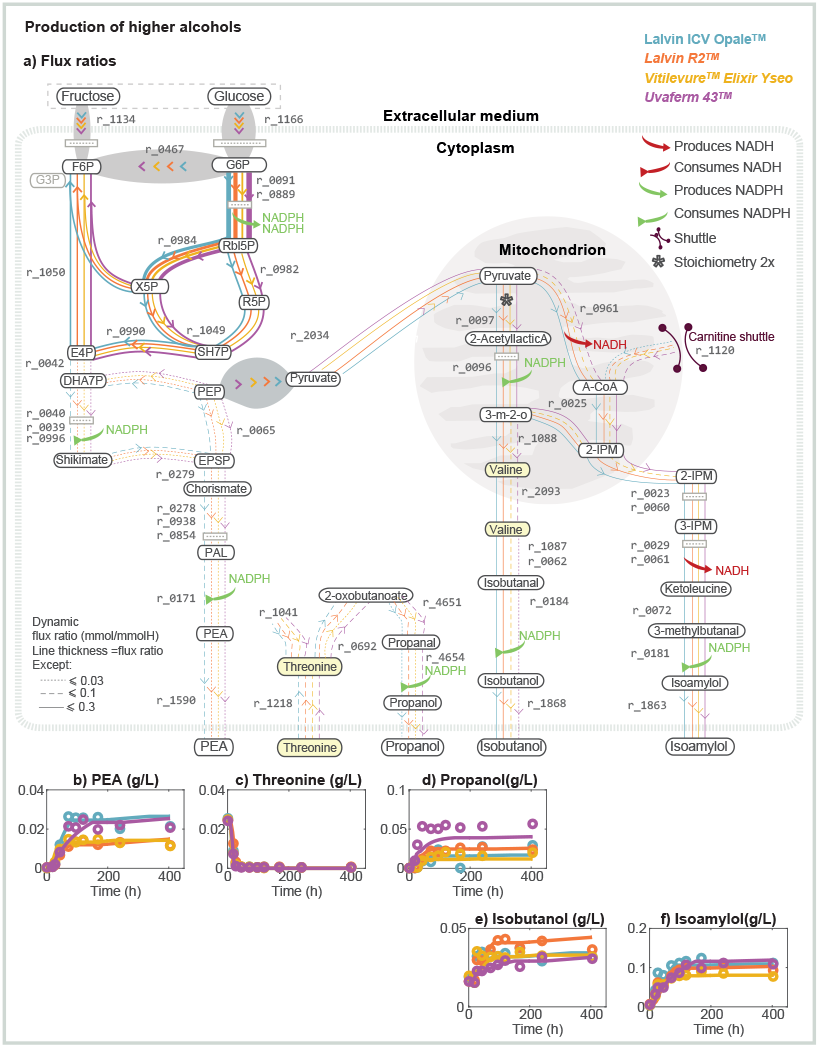
Overview of higher-alcohol production. Panel a depicts the predicted intracellular flux dynamic flux ratios related to higher alcohols: propanol, 2-phenylethanol (PEA), isobutanol, and isoamylol during the carbohydrate accumulation phase and their related effect on the redox balance of cofactors. Panels b to f represent a comparison between model predictions and experimental values of PEA, threonine, propanol, isobutanol, and isoamylol, respectively.

**Figure 4.**
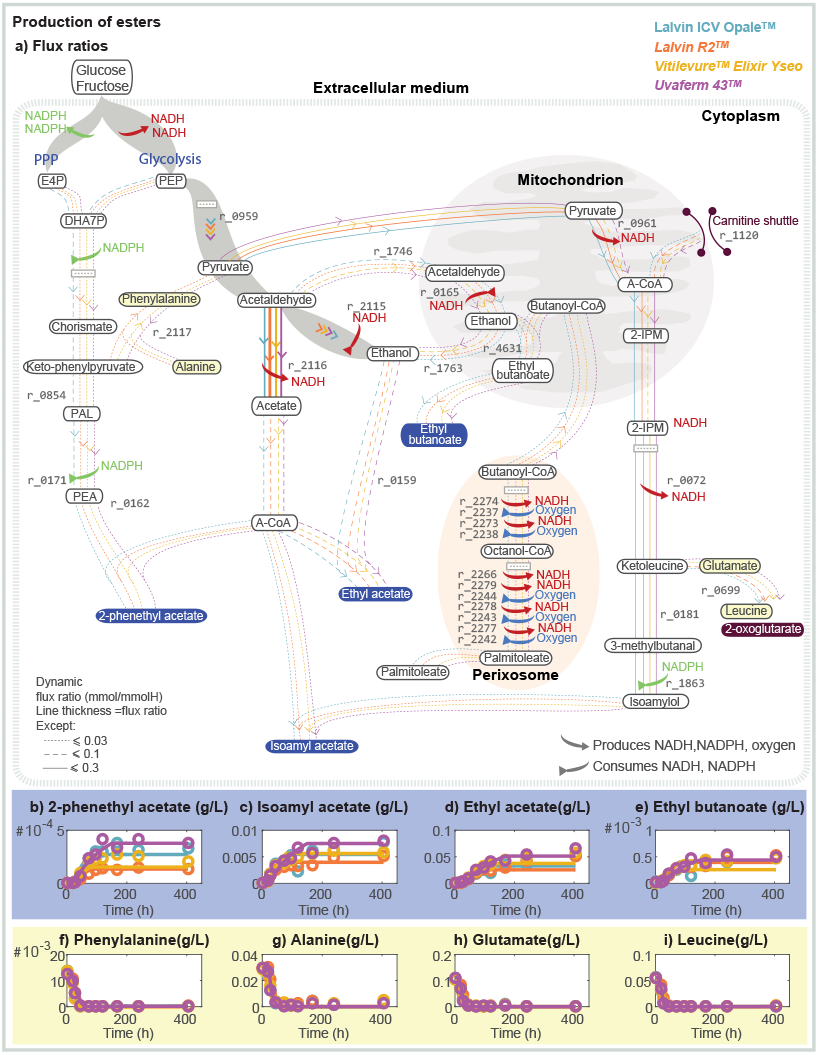
Overview of acetate and fatty acid ethyl ester production. Panel a shows the predicted intracellular dynamic flux ratios corresponding to the production of esters: 2-phenylethyl acetate, isoamyl acetate, ethyl acetate, and ethyl butanoate during the carbohydrate accumulation phase and their respective impact on the cofactor balance. Panels b to i correspond to the comparison between model predictions and experimental data of 2-phenylethyl acetate, isoamyl acetate, ethyl acetate, ethyl butanoate, respectively.

For the consumption of nitrogenous compounds, the model simulations obtained excellent fits against experimental data with each of the amino acid dynamic profiles containing a median R^2^ > 0.92 except for histidine (0.83), lysine (0.80), and asparagine (0.79). The model also simulated kinetic curves that were in reasonably good agreement with experimental data for the VOCs for most strains that have R^2^ > 0.85 (see Table S1). However, the VOCs of some strains (Opale and R2) e.g., ethyl hexanoate were predicted with an R^2^ = 0.22 and 0.16, respectively (Table S1 and Fig. S2). There was only a slight underprediction of propanol for the Uvaferm strain (see Fig. 3).

When examining the dynamic concentration profiles, the most apparent differences among the strains correspond to extracellular metabolites associated with core carbon, nitrogen, and lipid metabolism. These notable differences are evident when observing the secretion dynamics of acetate, glycerol, and succinate (Fig. 2) as well as the overall production of many VOCs, e.g., propanol, isoamylol, isobutanol, isoamyl acetate, phenylethyl acetate, and ethyl butanoate (Fig. 3 and Fig. 4).

### 3.3 Key flux predictions across various cell phases suggest the most distinct strain behavior occurs during the carbohydrate accumulation phase

The model framework was employed to elucidate the metabolic mechanisms adopted by the strains to achieve different compound concentration levels. Using the *Yeast 8.5.0* combined with the dFBA model framework, it is possible to predict the change of the entire set of fluxes throughout the simulated fermentation. The dynamic flux ratios were determined using the aforementioned equations (see Section 2.5 in Materials and Methods). Their values for some of the key intracellular fluxes between 6- and 40-hours during fermentation (carbohydrate accumulation phase) are presented in Supplementary Figure S1.

Essential fluxes concerning an anaerobic nitrogen-limited yeast phenotype were observed in all growth phases (see supplementary data files)‥ These fluxes pertained to specific pathways of the central carbon, nitrogen, and lipid metabolism. This key set of fluxes suggests the role of NAD-dependent acetaldehyde dehydrogenase (r_2115) to restoring redox balance and allow anaerobic growth at differing levels among the strains as suggested previously (Vargas et al., 2011, Scott et al., 2021b). The model predictions for growth were similar among the strains except for the Uvaferm strain, which experienced a later onset of stationary phase (150 h) compared to other strains (80 h) (Table S1). This distinction could be related to dissimilarity in protein turnover and synthesis especially pertaining to the Uvaferm strain.

The dynamic flux ratios pointed to aspartate-semialdehyde dehydrogenase (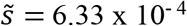, 1.05 × 10^−3^, 1.22 × 10^−4^, and 9.36 × 10^−4^ mmol/mmolH for Opale, R2, Elixir, and Uvaferm respectively), homoserine dehydrogenase (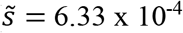, 1.05 × 10^−3^, 1.22 × 10^−4^, and 9.36 × 10^−4^ mmol/mmolH for Opale, R2, Elixir, and Uvaferm respectively), and glycerol-3-phosphate dehydrogenase (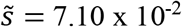, 8.00 × 10^−2^, 6.31 × 10^−2^, and 7.95 × 10^−2^ mmol/mmolH for Opale, R2, Elixir, and Uvaferm, respectively) (where mmolH is millimoles of consumed hexose) (shown as r_0219, r_0547, and r_0492 in Fig. 2) as the responsible for strain-dependent behavior. These differences relate to the variation in extracellular productions of glycerol and succinate (Fig. 2 a, d) The homoserine dehydrogenase enzyme is involved withing the aspartate where it regulates the NAD(P)-dependent reduction of aspartate beta-semialdehyde into homoserine which subsequently leads the biosynthesis of methionine, threonine, and isoleucine. In addition, the glycerol-3-phosphate dehydrogenase enzyme is responsible for catalyzing the NADH-aided reduction of dihydroxyacetone phosphate to glycerol 3-phosphate which leads to the biosynthesis of phospholipids.

As nitrogen limitation conditions arise, yeast metabolism circulates more sugar flux into the fermentation. As such, there is a decline in cell growth, and what is not spent (sugar, nitrogen, etc., flux) on biomass synthesis is directed to other routes. Overall, for Uvaferm, the decline in protein synthesis and lipid production lowered the demand for NADPH (see fluxes Table S2). At the onset of carbohydrate accumulation due to nitrogen limitation, restriction of NAD+ occurred. As nitrogen limits the rate of glucose and fructose uptake in conjunction with growth, the model opted to employ the glyoxylate cycle to reduce NADH production while still producing necessary tricarboxylic acid (TCA) cycle intermediates (Fig. 2). Overall, in this regard, the model simulations underscore the tight link between glycolysis, the TCA cycle, lipid, and amino acid metabolism under nitrogen-limited conditions.

Under nitrogen-limited fermentation conditions, sluggish fermentation can occur due to lower relative protein turnover, resulting in higher relative amounts of RNA and storage carbohydrates, e.g., trehalose (Varela et al., 2004). Furthermore, the release of C6 sugars from trehalose and glycogen mainly occurs during the onset of the stationary phase (Schulze et al., 1996). For the strains in this study excluding the Uvaferm, this release was predicted to occur at the carb. accumulation phase Uvaferm showed comparably lower flux values for both the carbohydrate pseudo-reaction 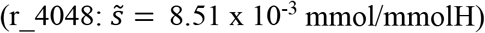 and trehalose-phosphatase 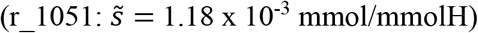. However, Uvaferm was the only strain to have predicted flux for both reactions during the stationary phase. This particularity suggests intracellular mechanisms bring about carbohydrate storage or accumulation for the other strains or later for Uvaferm. Furthermore, a later carbohydrate accumulation could affect the fermentative performance and capacity of the yeast strain (François and Parrou, 2001, Pérez-Torrado and Matallana, 2015).

Yeast cells produced most of the succinate during the carbon accumulation phase, where the Uvaferm strain contained a higher dynamic flux ratio than the other strains (Table S3, e.g., r_1000). This higher succinate concentration in Uvaferm is caused by cells taking up a preferred nitrogen source, i.e., glutamate via the GABA shunt which becomes deaminated by NAD+ dependent glutamate dehydrogenase to discharge α-ketoglutarate and ammonium (Fig. 2, r_0471). Consequently, the increase in the intracellular α-ketoglutarate concentration increases enzyme activities of the oxidative branch of the TCA cycle, causing succinate production. Previous works have alluded to the fact that carbon skeletons derived from deaminated glutamate are easily facilitated into the TCA cycle and, hence, transformed into succinate (Tesnière et al., 2015). This premise is supported by the model, which uses an aspartate transaminase-associated reaction (r_0216) to link glutamate and thus steer succinate formation (Fig. 3).

Although Uvaferm strain experienced substantial flux 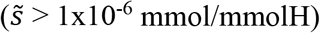 in many reactions associated with redox balance and acetyl CoA (e.g., r_2242, r_2266, r_2274, and r_2278, r_2279 as shown in Fig. 4) during the stationary phase while the other strains did not show any flux, the intracellular behavior differs among the strains with higher standard deviations (see supplementary data) most significantly during the carbohydrate accumulation phase. The exponential and carbohydrate accumulation phases also coincide with the vigorous production of higher alcohols and esters, desirable in fermented beverages such as wine and beer and commercially crucial for fragrances (for more info see Fig. S1 and S2). For instance, isoamyl acetate known for producing fruity aromas, experienced predicted 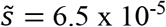, 5.3 × 10^−5^, 6.5 × 10^−5^, and 6.8 × 10^−5^ mmol/mmolH during the exponential and carbohydrate acc. phases whereas during the stationary phase the Uvaferm strain was the only strain to experience a flux which was 6.8 × 10^−5^ mmol/mmolH.

### 3.4 The role of acetate and ethyl esters in the transitions from growth to the stationary phase

Esters are a class of compounds that imbue a range of desirable fruity and floral aromas to wines and beer (Hirst and Richter, 2016). These esters are produced by a condensation reaction between acetyl or acyl-CoA and an alcohol (Saerens et al., 2010). Two types of esters are synthesized, acetate esters and fatty acid ethyl esters, respectively, based on whether acetyl-CoA or acyl-CoA is employed when forming them (Dzialo et al., 2017). More specifically, acetate esters are synthesized via the condensation of higher alcohols with acetyl-CoA, catalyzed by alcohol acetyltransferase enzymes (AATs) (Mason and Dufour, 2000). Acetate esters, primarily ethyl acetate, isoamyl acetate, isobutyl acetate, and 2-phenylethyl acetate, are quantitatively the most abundant type of esters formed during fermentation.

Our modeling analysis illustrated that acetate ester production was predominately modulated by the shifting of the acetyl-CoA/CoA ratios as the yeast cells transitioned from the exponential to the carbohydrate accumulation phase. This reflection was seen most strikingly from examining reactions associated with acetyl-CoA synthetase, alcohol acetyltransferase, malate synthase, and serine O-acetyltransferase (*ACS*s, *ATF*s, *MS*, and *SAT* related to reactions: r_0112, r_0113, r_0158 - r_0162, r_0716, and r_0992) where the strains that maintained the highest flux ratios between phases, produced the most acetate esters.

For instance, during the exponential phase, the model predicted a significant flux from isoamylol to isoamyl acetate inside the cytoplasm (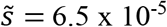, 5.3 × 10^−5^, 6.5 × 10^−5^, and 6.8 × 10^−5^ mmol/mmolH for Opale, R2, Elixir, and Uvaferm respectively; Fig. 4.; r_0160). This reaction sustained the same fluxes for all the strains during the carbohydrate accumulation phase. The same behavior was also observed in other ATF-associated reactions. In this reaction (r_0160), acetyl-CoA is consumed to form CoA, thereby illustrating that strains with higher flux ratios of acetyl-CoA/CoA were superior in producing higher acetate ester fluxes. This simulation result has been previously explored experimentally by Hong and co-workers (Hong et al., 2019). They showed that acetate ester formation in yeast is modulated by changing the CoA and acetyl-CoA levels by deleting and overexpressing the *BAP2* and *ATF1* genes, respectively (Hong et al., 2019). Furthermore, it has been shown that commercial yeast strains differ in the production of acetate esters, by orders of magnitude, even under identical fermentation conditions (Steensels et al., 2014). Our model and those previous results suggest that optimizing the activity of genes responsible for regulating acetyl-CoA/CoA could be beneficial in obtaining the appropriate acetate ester yield. For example, a researcher could adapt a platform tuning the enzyme expression of acetyl-CoA carboxylases for acetate esters, similar to a recent study for improving supplies of acetyl CoA and NADPH and eventual production of 3-hydroxypropionicacid (Qin et al., 2020).

Although there is much certainty regarding the mechanisms that form acetate esters during fermentation, the mechanisms, including genetics and regulation of fatty acid ester formation, are still less clear. However, medium-chain fatty acids (MCFA) intermediates are prematurely released from the cytoplasmic fatty acid synthase (FAS) complex during the late exponential growth phase. This release triggers ester synthesis as pointed out by (Taylor and Kirsop, 1977). Subsequently, CoA can activate an MCFA in combination with ATP and ethanol to enzymatically form an MCFA-ethyl ester (Saerens et al., 2010). This mechanism can be observed in our predictions in Fig. 4 (e.g., r_4631). Additionally, transitions in fluxes were noticed in FAS reactions (r_2140 and r_2141, fatty-acyl-CoA synthase) from the exponential growth phase to the carbohydrate accumulation phase. For instance, these shifts in flux values among the strain between growth phases could correspond to the activity in CoA with MCFA. This elaborate relationship causes the levels of fatty acid esters to depend significantly on lipid metabolism and acetyl-CoA.

Several investigations indicate at least three modulating routes for the flow of MCFAs to produce ethyl esters. Firstly, the upregulation of fatty acid synthases (*FAS1* and *FAS2*) and acyltransferases (*EEB1* and *EHT1*) (Saerens et al., 2010). Secondly, reduced acetyl-CoA carboxylase activity has been shown to play a role where the inhibition of acetyl-CoA carboxylase facilitates the discharge of MCFAs from the *FAS* complex (Hirst and Richter, 2016). Thirdly, an increase in MCFA concentrations induces higher concentrations of ethyl esters in wine (Saerens et al., 2008).

Here, it was inferred the model is in line with the hypothesis that these three routes play a significant factor in producing MCFA ethyl esters. Higher fluxes during the exponential and carbohydrate accumulation phases in reactions associated with fatty-acyl-CoA synthase (n-C16:0CoA), fatty-acyl-CoA synthase (n-C18:0CoA), alcohol acyltransferase (butyryl-CoA), and alcohol acyltransferase (hexanoyl-CoA) (r_2140, r_2141, r_4631, and r_4629) resulted in the higher overall production of ethyl butanoate and ethyl hexanoate – essential MCFA ethyl esters found in wines - for the Uvaferm and Opale strains (Fig. 4 and 5). For instance, Uvaferm and Opale experienced 71.1% and 12.6%, respectively, more flux than the Elixir strain for the reaction associated with fatty-acyl-CoA synthase (n-C16:0CoA), which resulted in 14.4% and 8.2% more overall production for Uvaferm and Opale, respectively, than the Elixir strain. Conversely, when examining the effect of acetyl-CoA carboxylase, the strain (R2) that experienced the lowest flux value for an acetyl-CoA carboxylase-associated reaction (r_0109) during the exponential phase showed the highest flux for the formation of ethyl butanoate during the carbohydrate accumulation phase (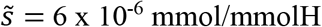, r_4646).

**Figure 5.**
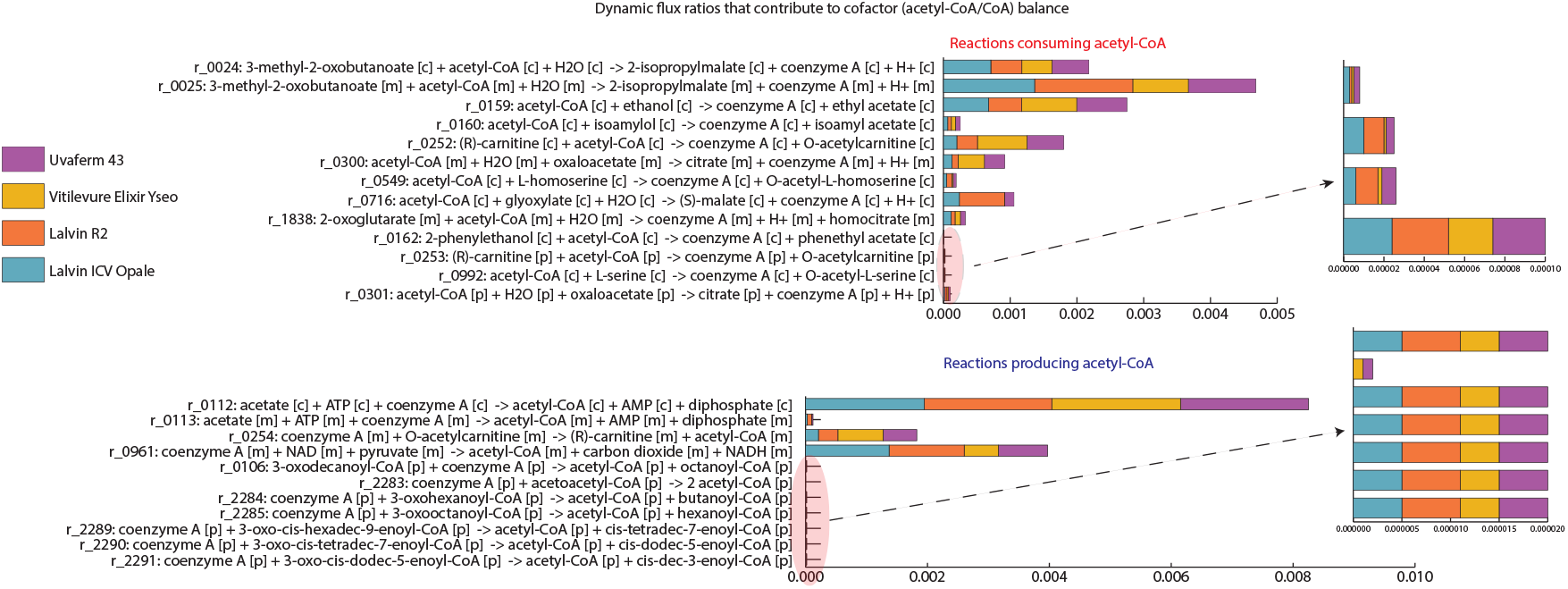
Comparative illustration of the fluxes through the reactions consuming and producing acetyl-CoA during the carbohydrate accumulation phase, illustrating how acetyl-CoA/CoA ratios modulate cellular phenotype.

Lastly, a relationship was deduced by observing the intracellular fluxes related to the production and consumption of CoA (Fig, 5). For instance, we could see an uptick or higher flux in ethyl esters for the strains that experienced more significant imbalances of CoA and acetyl-CoA (based on greater flux values in the direction of acetyl-CoA consumption). After evaluating the difference in CoA and acetyl-CoA producing/consuming reactions, it was determined that the Uvaferm strain had the most significant disparity. This result aligns with the prediction of the Uvaferm strain producing the most ethyl butanoate (Fig. 4). This detected mechanism for yeast to produce MCFA ethyl esters to correct imbalances of CoA and acetyl-CoA has been hypothesized before (Lambrechts and Pretorius, 2000). However, here we show that metabolic modeling supports this view. Also, since MCFAs are toxic to yeast and, at sufficient concentrations, can lead to stuck or sluggish fermentations (Viegas et al., 1989), strains better at producing ethyl esters from CoA are predicted to achieve a fitness advantage.

Taken together, we report a model able to predict the main routes of ester formation over the course of wine fermentation for different commercial yeast strains. Our results reveal that that ester formation depends on intracellular acetyl-CoA/CoA levels. In addition, we gained novel insights about metabolic routes taken to form key esters which pave the way for future metabolic engineering strategies to manage ester content in alcoholic fermentations. For instance, one can genetically modify an available GEM to delete, add, sub-express, or over-express, for a metabolite of interest (Copeland et al., 2012). Then, appropriately calibrate the model using experimental data, especially from sugar and nitrogenous substrates. Finally, one can examine the effect of genetic modification and/or varying the substrate composition on the predictions of certain aroma profiles.

## 4 CONCLUSION

As yeast becomes increasingly utilized by the food, biotechnology, and cosmetic sectors to produce desirable aroma compounds organically, it is crucial to understand how the most prominent of these alluring aromas are produced under industrially-related settings. By applying an adapted dynamic flux balance analysis framework, this study was able to predict many aspects of an experimental enological fermentation conducted using commercial wine strains, including the production of primary metabolites and biomass, as well as key aromas, e.g., esters. By observing from simulations that pivotal intracellular flux routes varied among the strains chiefly during the carbohydrate accumulation phase, underlying mechanisms related to redox balance and the utilization of acetyl-CoA were understood to be responsible for observed phenotypic behavior. Moreover, the dynamic genome-scale modeling approach allowed the study of individual fatty acid fluxes over time, supporting the potential role that certain ethyl esters form to remove toxic fatty acids from the cell by previous experimental work. Future research should look further into the genes regulating the acetyl-CoA, as mentioned above, with other commercial strains. This type of research could establish a clearer, more general picture of how various types of aromas are produced during yeast fermentation. Overall, the predictions agreed with previous experimental findings and illustrated which metabolic pathways play a role in aroma production, making this a promising approach for future use in studies related to individual fluxes of important metabolites in oenological conditions and for comparing metabolic differences between commercial wine yeast strains.

## ACKNOWLEDGEMENTS

E. B-C. acknowledges funding from MCIU/AEI/FEDER, UE (grant reference: PID2021-126380OB-C32), and Xunta de Galicia (IN607B 2020/03). In addition, some materials and resources were funded by a Wageningen University and Research internal fund from the Food Microbiology department.

## Authors’ contributions

### CRediT (Contribution Roles Taxonomy)

Conceptualization, W.T.S.J., D.H, E.J.S., R.A.N., and E.B-C.; Data Curation, W.T.S.J. and D.H., Formal Analysis, W.T.S.J., D. H., and E.B-C.; Funding Acquisition, E.J.S. and E.B-C; Investigation, W.T.S.J.; Methodology, W.T.S.J., D.H., and E.B-C.; Resources, R.A.N, and E.B-C.; Software, D.H, and E.B-C.; Supervision, E.J.S, R.A.N., and E.B-C.; Validation, W.T.S.J.; Visualization, W.T.S.J., D.H., and E. B-C; Writing—original draft, W.T.S.J.; Writing—review and editing, W.T.S.J., D.H., E.J.S., R.A.N., and E.B-C. All authors have read and agreed to the published version of the manuscript.

## CONFLICT OF INTEREST

The authors declare no competing interests

## Notes

### Competing Interest Statement

The authors have declared no competing interest.

### Summary of Updates

Results and Discussion Section updated

