## Supplementary material for "Dynamic genome-scale modeling of *Saccharomyces cerevisiae* unravels mechanisms for ester formation during alcoholic fermentation": Table S1

Table S1. R2 values for experimental data vs. model predictions among the yeast strains for each metabolite and biomass

| | uvaferm | R2 | opale | elixir | Median |
|---|---|---|---|---|---|
| biomassX_o | 0.9862 | 0.965 | 0.9572 | 0.9086 | 0.9611 |
| glucose_o | 0.994 | 0.9865 | 0.9643 | 0.9293 | 0.9754 |
| fructose_o | 0.9728 | 0.9523 | 0.9538 | 0.9543 | 0.95405 |
| glycerol_o | 0.9855 | 0.9508 | 0.327 | 0.8264 | 0.8886 |
| ethanol_o | 0.9987 | 0.9917 | 0.5427 | 0.9694 | 0.98055 |
| succinate_o | 0.8288 | 0.9729 | 0.4766 | 0.7861 | 0.80745 |
| acetate_o | 0.62 | 0.9514 | 0.2702 | 0.8925 | 0.75625 |
| ethylacetate_o | 0.9214 | 0.3555 | 0.7104 | 0.7833 | 0.74685 |
| isobutylacetate_ | 0.4737 | 0.7337 | 0.6094 | 0.7713 | 0.67155 |
| isobutanol_o | 0.8214 | 0.8467 | 0.0225 | 0.087 | 0.4542 |
| isoamylacetate_ | 0.9293 | 0.7989 | 0.6084 | 0.9165 | 0.8577 |
| isomaylalcohol_ | 0.9537 | 0.8482 | 0.7797 | 0.9264 | 0.8873 |
| phenylacetate_o | 0.9309 | 0.9519 | 0.7893 | 0.2547 | 0.8601 |
| phenylethanol_o | 0.847 | 0.8764 | 0.8984 | 0.7878 | 0.8617 |
| histidine_o | 0.9457 | 0.7882 | 0.8752 | 0.7751 | 0.8317 |
| glycine_o | 0.9826 | 0.99 | 0.9829 | 0.9668 | 0.98275 |
| alanine_o | 0.9681 | 0.9298 | 0.9572 | 0.9378 | 0.9475 |
| glutamate_o | 0.9922 | 0.94 | 0.9836 | 0.9554 | 0.9695 |
| glutamine_o | 0.9947 | 0.9786 | 0.9848 | 0.9253 | 0.9817 |
| isoleucine_o | 0.9109 | 0.9314 | 0.99 | 0.9859 | 0.95865 |
| aspartate_o | 0.9226 | 0.8964 | 0.9833 | 0.9649 | 0.94375 |
| arginine_o | 0.9925 | 0.9799 | 0.9708 | 0.9231 | 0.97535 |
| asparagine_o | | | 0.803 | 0.7683 | 0.78565 |
| serine_o | 0.8949 | 0.8999 | 0.9783 | 0.9667 | 0.9333 |
| tyrosine_o | 0.9765 | 0.9452 | 0.9819 | 0.9625 | 0.9695 |
| tryptophan_o | 0.9778 | 0.9806 | 0.9402 | 0.9218 | 0.959 |
| threonine_o | 0.9807 | 0.989 | 0.8979 | 0.9165 | 0.9486 |
| leucine_o | 0.9978 | 0.9454 | 0.997 | 0.968 | 0.9825 |
| valine_o | 0.9867 | 0.9818 | 0.9344 | 0.801 | 0.9581 |
| phenylalanine_o | 0.9497 | 0.9673 | 0.9921 | 0.9989 | 0.9797 |
| lysine_o | 0.7259 | 0.7058 | 0.8823 | 0.936 | 0.8041 |
| methionine_o | 0.8767 | 0.9039 | 0.9663 | 0.9406 | 0.92225 |
| ammonia_o | 0.7024 | 0.6837 | 0.9518 | 0.9259 | 0.81415 |
| ethylhexanoate_ | 0.6288 | 0.1602 | 0.2147 | 0.5722 | 0.39345 |
| propanol_o | 0.2842 | 0.8483 | | | 0.56625 |
| ethyl_butanoate_ | 0.9028 | 0.7986 | 0.546 | 0.2568 | 0.6723 |
