## Supplementary material for "Dynamic genome-scale modeling of *Saccharomyces cerevisiae* unravels mechanisms for ester formation during alcoholic fermentation": Figure S1

A)

| | opale | R2 | elixir | uvaferm |
|---|---|---|---|---|
| LAG | 0 | 0 | 0 | 0 |
| exponentia | 5.98E+00 | 6.73E+00 | 6.21E+00 | 7 |
| carb accum | 3.38E+01 | 3.68E+01 | 3.66E+01 | 37 |
| Stationary | 8.00E+01 | 7.97E+01 | 8.00E+01 | 150 |
| Decay | 1.00E+02 | 1.00E+02 | 1.00E+02 | 170 |
| Final | end | end | end | end |

$$S_{i,G} = \frac{\int_{t_L}^{t_S} v_i(t) \cdot \mathrm{DW}(t)}{\int_{t_L}^{t_S} v_{\mathrm{Glx}}(t) \cdot \mathrm{DW}(t) + \int_{t_L}^{t_S} v_{\mathrm{Fr}}(t) \cdot \mathrm{DW}(t)}$$

B)

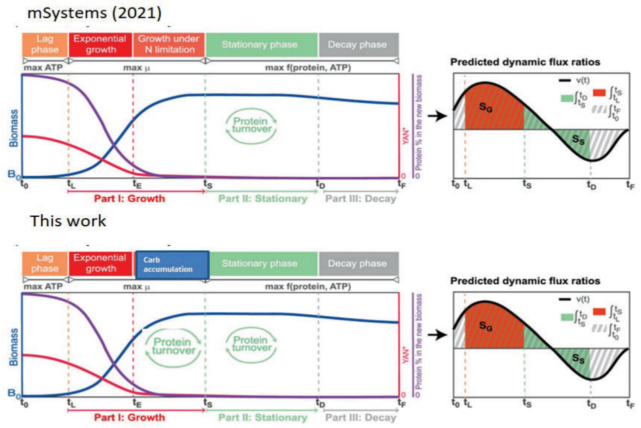


C)

In each phase, we computed the integral of each flux multiplied by the biomass over time and normalized its value with the accumulated flux of consumed hexoses (glucose and fructose).

```
vG=trapz(TSIM(time_index),scale(findRxnIDs(model,index_G),time_index));
vF=trapz(TSIM(time_index),scale(findRxnIDs(model,index_F),time_index));

for i=1:(size(scale,1)-2)

    AUC(i)=trapz(TSIM(time_index),scale(i,time_index));

%     if abs(-AUC(i)/(vG+vF) )>0.00005
        form=printRxnFormula(model,'rxnAbbrList',model.rxns{i},'metNameFlag',true);
        fprintf(fid,'%f\t%s\t%s\t%f\t%f\t%s\n',-AUC(i)/(vG+vF),model.rxnNames{i},model.rxns{i},model.lb(i),model.ub(i),form{1});
%     end
end
```

Figure S1. A) Illustrating the simulated duration of the cell phases as well as B) giving a more detailed outline of the similarities and differences between the modeling approach used in this study vs. Henriques et. al .2021. C) A depiction of script used for computing the dynamic flux ratios based on normalizing the carbohydrate. accumulation.
