## Supplementary material for "Dynamic genome-scale modeling of *Saccharomyces cerevisiae* unravels mechanisms for ester formation during alcoholic fermentation": Figure S2

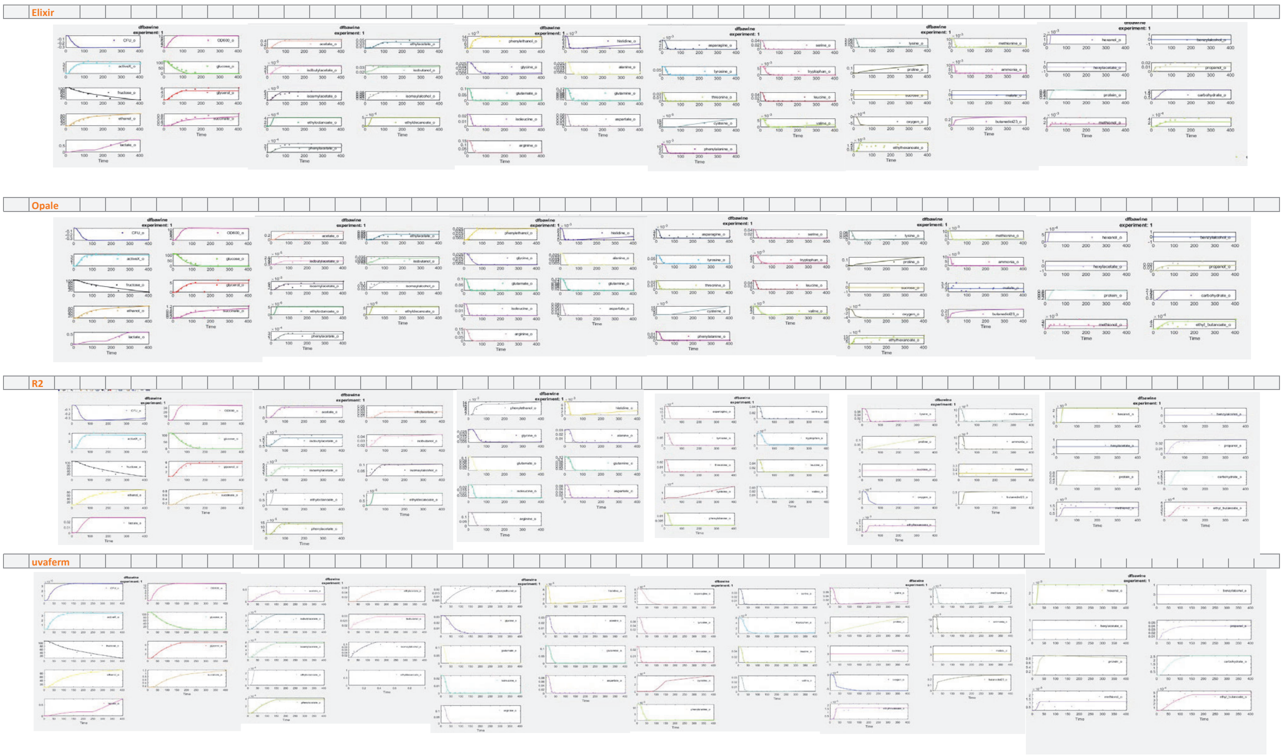


Figure S2. Kinetic curves showing the predicted concentrations/biomass against experimental data over time for all the metabolites of interest for each strain.
